# Wave-like SnRK1 Activation and Tre6P–Sucrose Imbalance Shape Early Salt Stress Response in Growing Alfalfa Leaves

**DOI:** 10.1101/2025.04.23.650249

**Authors:** Barbieri Giuliano, Parola Rodrigo, Feil Regina, Rodriguez Marianela S.

## Abstract

Alfalfa (*Medicago sativa* L.) is a key forage crop valued for its adaptability and nutritional quality, yet salinity significantly limits its productivity, particularly in arid regions. Understanding early stress responses is crucial for improving resilience. Salt stress impairs leaf growth and photosynthesis, triggering complex, time-dependent signaling. Sucrose non-fermenting kinase 1 (SnRK1), a central metabolic sensor, regulates metabolism and growth under stress. We investigated the dynamics of SnRK1, sucrose, and trehalose-6- phosphate (Tre6P) during leaf expansion in a salt-tolerant alfalfa cultivar. Plants were hydroponically grown and exposed to 200 mM NaCl. Stress induced transient declines in chloroplast performance (Fv/Fm, performance index). SnRK1 activity peaked within 1 hour post-treatment (hpt), likely initiating metabolic shifts. By 3 hpt, sugar metabolism shifted, with increased catabolism, TCA cycle modulation, and glucose-6-phosphate accumulation. SnRK1 and sucrose showed opposing wave-like patterns, with sucrose peaking at 1 day post-treatment (dpt) as Tre6P declined, indicating a disrupted regulatory link. A second SnRK1 peak at 3 dpt correlated with reduced leaf growth. Exogenous sucrose inhibited SnRK1, while NaCl enhanced it. This is the first report of wave-like SnRK1 activation and Tre6P–sucrose uncoupling in alfalfa, highlighting early SnRK1 activation as key to salt stress adaptation.

**Highlights:**

1. Early SnRK1 activation is a key determinant of salt stress response in alfalfa, linking early biochemical shifts to downstream metabolic alteration.
2. First evidence of a wave-like SnRK1 activation pattern and disruption of the Tre6P– Sucrose nexus reveals novel dynamics in alfalfa’s stress signaling.
3. Sucrose inhibits SnRK1 activity in source leaves with or without NaCl

## Introduction

Alfalfa (*Medicago sativa L.*) is the most important perennial forage legume crop playing a key role in meat and milk production worldwide and providing vital ecosystem services (Ciacci *et al*., 2023; Basigalup, 2022). Its abilities to fix nitrogen, sequester carbon, and improve soil structures highlight its role in enhancing agricultural sustainability (Ciacci *et al*., 2023). However, soil salinization represents a significant challenge to crop productivity, affecting germination, growth, and biomass accumulation. Globally, over 6% of the land area is impacted by salinity or sodicity, with millions of hectares experiencing declining productivity or being removed from cultivation annually (FAO and ITPS, 2015). Alfalfa suffers biomass and yields reductions of up to 35% under saline conditions (Al-Farsi *et al*., 2020). The genetic improvement of alfalfa’s salt tolerance remains challenging due to its complex physiological and genetic responses, involving multiple genes and biochemical mechanisms.

Salinity stress negatively impacts leaf growth while the photosynthetic processes are affected by induced stomatal closure, inhibition of chlorophyll biosynthesis and damage of the photosynthetic apparatus, disrupting electron flow between photosystems II (PSII) and I (PSI) (Qin *et al*., 2021). Chlorophyll fluorescence (ChlF) transients provide valuable insights into the electron transport components within PSII and PSI (Bussotti, 2010). The OJIP curve is particularly useful for analyzing how environmental stresses affect the electron transport chain (Kalaji *et al*., 2014). Under stress conditions, additional K and L bands offer further information on electron flow from the oxygen-evolving complex (OEC) to PSII and connectivity between PSII units, respectively (Oukarroum *et al*., 2007; Strasser *et al*., 2004).

The primary output of photosynthesis is sugar. Therefore, since the photosynthetic process is negatively affected by salt stress, the synthesis of sugars may be reduced. However, plants under salt stress tend to accumulate sugars, indicating an alteration in carbon utilization, sink strength, growth rate, and stress response (Henry *et al*., 2015). The inability to efficiently manage carbon resources has clear penalties for plant growth. In this regard, the sugar accumulation in leaves may impact negatively on photosynthesis as a feedback negative mechanism (Rodriguez *et al*., 2010).

Plant sugars are essential for energy provision, osmotic balance, and signaling, with sucrose (Suc) homeostasis being vital for optimal growth and development under adverse conditions (Ruan, 2014; Pereyra *et al*., 2023). In the proposed model called Suc-trehalose- 6-phosphate (Tre6P) nexus, Suc levels in plants are strongly correlated with Tre6P in normal conditions, which links growth and development to their metabolic status across various environmental scenarios (Yadav *et al*, 2014; Fichtner and Lunn, 2021). Although Suc accumulation in salt-stressed alfalfa leaves has been well-documented (Gao *et al*., 2018), the underlying mechanisms remain poorly understood, and there is a lack of information on how the components regulating carbon assimilation, homeostasis and utilization for growth are coordinated. Furthermore, Tre6P inhibits sucrose non-fermenting-related protein kinase (SnRK1) in sink tissues (Zhang *et al*., 2009).

SnRK1 is a hub kinase which regulates energy homeostasis and stress responses in plants through transcriptional, translational, and metabolic reprogramming, inducing catabolic pathways and suppressing anabolic pathways (Baena Gonzalez *et al*., 2007; Polge and Thomas, 2006; Ramon *et al*., 2019). When subjected to saline stress, plants alter their energy expenditure and slow down their growth to adapt to stress conditions (Hartmann *et al*., 2015). Energy homeostasis is a key strategy for plant salinity tolerance, which could provide valuable tools for breeding programs (Tyerman *et al*., 2018). In this context, SnRK1 has a leading role inducing genes related to nutrient remobilization and gluconeogenesis, while repressing genes related to protein synthesis, the tricarboxylic acid (TCA) cycle, and glycolysis (Peixoto and Baena Gonzalez, 2022). A positive correlation between SnRK1 activation and salt-tolerance was demonstrated by numerous studies (Barajas-Lopez *et al*., 2017; Rodriguez *et al*., 2019; Wang *et al*., 2020). However, the role of SnRK1 in salt stress has primarily been studied in a specific time without distinguishing phase-specific responses, leaving significant gaps in understanding its dynamic activity in leaves, particularly in alfalfa.

Plant responses to salinity unfold in two distinct phases. The initial phase occurs rapidly, ranging from minutes to days, and is associated with sodium detection and signaling (Roy *et al*., 2014). During this early stage, osmotic stress hampers water absorption, while prolonged exposure leads to ion toxicity (Munns and Tester, 2008). During early salt stress stages, SnRK1 could adjust gene expression by activating vital transcription factors (e.g. ABI5, bZIP63), essential for ABA-mediated stress responses and energy metabolism (Baena González *et al*., 2007). Further studies in Arabidopsis roots have suggested that SnRK1 supports stress responses by facilitating significant metabolic reprogramming through the coordinated activation of bZIP transcription factors during salt stress (Hartmann *et al*., 2015). With prolonged salt stress, SnRK1 could fulfill a different role, inducing plant resilience through cellular structure stabilization and ionic balance maintenance.

Given that leaves account for over 70% of alfalfa’s total protein yield (Hojilla-Evangelista *et al*., 2016), unraveling the impact of salinity on leaf development and metabolic pathways is critical for sustaining crop quality and productivity. Despite its key role in metabolic regulation, SnRK1 remains underexplored in the context of time-dependent metabolic changes in alfalfa, with no studies examining its dynamics during salt stress. This study aims to bridge this gap by providing detailed insights into the temporal interplay of leaf growth, photosynthesis, metabolic alterations, and SnRK1 activity in alfalfa under salt stress. Furthermore, the underexplored interaction between Suc, Tre6P, and SnRK1 during salt stress offers a unique opportunity to uncover novel regulatory mechanisms in alfalfa.

This study aimed to: (1) investigate the time-dependent effects of salt stress on leaf expansion, photosynthesis and metabolic alterations in alfalfa leaves, (2) analyze the Suc- Tre6P-SnRK1 signaling network under salinity, and (3) provide the first dynamic analysis of SnRK1 activity in alfalfa leaves. We hypothesize that SnRK1 rapidly responds to salt stress, triggering metabolic reprogramming and adaptive mechanisms to enhance tolerance.

Our findings reveal the time-dynamic changes in SnRK1 activity in alfalfa, underscoring its pivotal role in early and sustained stress responses. By integrating physiological, biochemical, and molecular analyses, this study provides a comprehensive understanding of how alfalfa leaves adapt to salinity, offering insights into key pathways that could inform breeding strategies for enhanced salt tolerance.

## Materials and Methods

### Biological Material and experimental setup

Alfalfa seeds of the salt-tolerant Argentinian cultivar Kumen PV INTA (Pecetti *et al*., 2024), a non-dormant with a rest grade of 8 and resistant to Phytophthora, anthracnose, and fusarium wilt, were provided by the alfalfa breeding program of the National Institute of Agriculture Technology (INTA).

Seeds were germinated on mesh supports with moist vermiculite placed over 3-liter containers with 0.5X B&D medium (Broughton and Dilworth, 1971) and continuous aeration. The Kumen cultivar used in the study was chosen due to its high biomass production, relative salt tolerance, and Na^+^ and Cl^−^ exclusion traits under saline conditions in field trials (Cornacchione and Suarez, 2016) and notable resilience and adaptive responses to salt stress shown in hydroponics (Barbieri *et al*., 2024). Three independent trials were carried out under controlled light (300 PAR) under a 16 h light/8 h dark cycle and temperature (23 ± 2°C) conditions in a growth chamber. The saline treatment (200 mM NaCl) was applied 20 days after planting. Sampling was performed on the third trifoliate expanded leaf (from top). Twelve plants per treatment were sampled at specific time points post-treatment 0; 1 and 3 hour post treatment (hpt); 1; 3- and 7-days post-treatment (dpt), samples were grouped into three pools of four leaves each (Figure S1).

### Effects of Sucrose and NaCl on SnRK1 activity in isolated leaves

Detached leaves of third trifoliate expanded leaves (from top) from 20 days-old plants grown in the same condition as mentioned above were incubated for 3 hours in light in Petri dishes containing 30 ml of the following solutions: control (H_2_O), 10 mM sucrose, 100 mM NaCl, and a combination of 10 mM sucrose + 100 mM NaCl. 24 h-dark treatment was used as SnRK1 activity positive control. After incubation, the leaves were ground to a fine powder using liquid nitrogen. Aliquots of 100 mg from the frozen powder were taken for extraction and analysis of SnRK1 activity, sodium and soluble sugars content.

### Leaf area and absolute leaf area growth rate (ALAGR)

To determine leaf area individual leaves were outlined, and leaf area was calculated using ImageJ software. The increase in leaf area over a specific period (absolute leaf area growth rate, ALAGR) was calculated as: ALAGR=A2−A1/t2−t1

Where:

A1: Leaf area at t1 (cm^2^)
A2: Leaf area at t2 (cm^2^)
t2−t1: Time interval for leaf growth measurement (days).

### Ion concentration

Fresh leaves (100 mg) were finely ground and dried at 70°C until completely desiccated. For extraction, each dry sample was resuspended in 1 ml of 30% CH_3_OH (v/v) prepared with Milli-Q water. The suspension was vortexed at room temperature for 15 min. Samples were centrifuged at 10000 rpm for 10 min, and 1 ml of the resulting supernatant was collected. To ensure purity, the supernatant was filtered through Nylon filters before proceeding with further analyses. Ions were determined by HPLC (Shimadzu Prominence Modular HPLC, Kyoto, Japan). A filtered and degassed 3 mM oxalic acid solution was used as eluent. Cations were determined using a Shim-pack IC-C3 (Shimadzu Co.) column with IC-C3G guard column. Each run was performed at 40 °C. flow rate was 1.2 ml/min, during 18 min, without suppression. The detection was performed by conductometry. Chromatography was registered and analyzed using LabSolutions (ver.5.6) software. (Pantsar-Kallio and Manninen, 1995).

### Chl fluorescence measurement on alfalfa leaves

The photosynthetic efficiency of each experimental group was determined with two ChlF methods: a) Pulse-amplitude modulated (PAM) handheld MultispeQ V2.0 device linked to the PhotosynQ platform (Kuhlgert *et al*. 2016; www.photosynq.org). The following parameters were analyzed at the third fully expanded leaves: maximal photochemical efficiency of PSII (Fv/Fm), relative electron transport rate (Phi2), chlorophyll content (SPAD), linear electron flow (LEF), number of opened PSII reaction centers(qL), proton conductivity (gH+), steady state proton flux (vH+), non-photochemical fluorescence quenching (PhiNO), non-photochemical quenching efficiency (PhiNPQ). The protocol is available on the PhotosynQ platform under the name Photosynthesis Rides 500 (https://photosynq.org). Measurements were taken once between 12 p.m. and 2 p.m. The parameters are described in Table S1.

b) Chlorophyll-a fluorescence transient (ChlF) was measured using a Pocket-PEA fluorometer (Plant Efficiency Analyzer, Hansatech Instruments Ltd., King’s Lynn, Norfolk, UK). The polyphasic OJIP fluorescence kinetics were utilized to assess PSII activity in both salt-treated and control alfalfa plants. Prior to measuring ChlF, the third fully expanded leaves were dark-adapted with leaf clips for at least 30 minutes to ensure full oxidation of the reaction centers (RC), achieving the minimum fluorescence (Fo or O step) at approximately 50 µs. A 1-second actinic light pulse of 3500 μmol photons m^−2^s^−1^ was then applied to reach the maximum fluorescence emission (Fm or P step) at around 300 ms. Intermediate steps, known as J and I, were recorded at 2 ms and 30 ms, respectively. The OJIP parameters were calculated using the software provided by the Pocket-PEA manufacturer, following the method described by Strasser *et al*. (2004).

Changes in the activity of the OEC and the connectivity at PSII were assessed following the methodology of Chen *et al*. (2015). To detect changes in the OEC (K-band), fluorescence curves were normalized from the O to the J step, resulting in WOJ curves. The WOJ curves from treated plants were then subtracted from those of the control plants ((WOJ_(Treated)_ - WOJ_(Control)_) to reveal the K bands (ΔWOJ). Positive values in these bands indicate damage to the OEC. For detecting L bands, curves were normalized between 0 and 300 μs and plotted as WOK curves. The difference between WOK_(Treated)_ and WOK_(Control)_ was used to generate ΔWOJ curves, which made the L bands visible. Positive values in the WOJ curves correlate with the loss of connectivity at PSII.

### Primary metabolite quantification

Alfalfa leaves were ground to a fine powder in liquid nitrogen and aliquots (15–20 mg) of frozen powder were extracted with chloroform/methanol as described by Lunn *et al*. (2006). Primary metabolites were measured by LC-MS/MS as described by Lunn *et al*. (2006) with modifications as described by Figueroa *et al*. (2015).

### Soluble Sugars

Frozen leaves samples (100 mg fresh weight) were ground to a fine powder using liquid nitrogen and homogenized in 1 ml of 80 % ethanol 4°C. The suspension was incubated at 30 °C for 30 min and then centrifuged at 12000 g for 10 min. The resulting extract was purified by adding 1 ml of pure acetonitrile, followed by a 30 min incubation at room temperature. The mixture was centrifuged again at 12000 g for 10 min. Subsequently, 1 ml of the supernatant was evaporated at 90 °C for 1h. Finally, the dried residue was resuspended in 700 μL of 75% acetonitrile and filtered using cellulose filters. Suc, Glc, fructose levels were determined using HPLC (Shimadzu) under an isocratic acetonitrile: water (81:19) flow (1 ml. min^-1^) at 30 °C. An amine column ZORBAX (analytical 4.6 x 250 mm, 5-micron, Agilent, USA) was used and the sugars were identified by their retention times using a RI detector and quantified using standards.

### SnRK1 activity determination

100 mg of each fresh leaves sample harvested after the salt treatment, was ground with mortar and pestle in presence of 500 µl cold extraction buffer containing 25 mM NaF, 100 mM Tricine-NaOH (pH 8.0), 2 mM tetrasodium pyrophosphate, 5 mM dithiothreitol, 0.5 mM ethylene glycol tetra acetic acid, 0.5 mM ethylene diamine tetra acetic acid, 1 mM phenylmethylsulfonyl fluoride, 1 mM benzamidine, 1 mM protease inhibitor cocktail (Sigma P9599), phosphatase inhibitors (PhosStop; Roche), and 2% (w/v) insoluble polyvinylpyrrolidone. The homogenate was transferred to two cold microfuge tubes and centrifuged at 12000 g for 5 min at 4 °C. The supernatant was desalted on a pre-equilibrated 2.5 ml centrifuge column (Sephadex G-25 medium columns; GE Healthcare). SnRKl activity was determined by the Universal Kinase Activity Kit (R&D Systems, Minneapolis, MN, United States, EA004) by using AMARA (AMARAASAAALARRR) (Sugden *et al*., 1999) polypeptide as the substrate.

### Immunoblot of SnRK1 T-loop phosphorylation

Phosphorylation at the alpha subunit conserved activation T-loop threonine was detected with a polyclonal antibody against the phospho-Thr-172 (P-Thr-172) present in the T-loop of AMPK, (Cell Signaling, Danvers, MA, USA). For immunoblot analysis, 20 μg of soluble protein were run and separated in 12% SDS-PAGE gels. Proteins were transferred onto polyvinylidene fluoride (PVDF) membranes and detected using secondary antibodies conjugated to alkaline phosphatase and revealed with an NBT (nitroblue tetrazolium chloride)/BCIP (5-bromo-4-chloro-3’-indolylphosphate p-toluidine salt) (Invitrogen catalog #34042).

### RNA isolation, cDNA synthesis, and qRT-PCR analysis

For gene expression analysis, pools of leaves from three alfalfa plants were used . Total RNA was extracted using the TRIZOL method. 2 µg of RNA DNA-free was used with oligo(dT) and random primers (Promega) for cDNA synthesis using M-MLV reverse transcriptase (Promega). The qPCR assays were performed in the iQ5 thermocycler (BioRad) with iQ SYBR Green Supermix (BioRad). Both qPCR and cDNA were performed according to the manufacturer’s instructions and expression levels were analyzed using the 2^-ΔΔCt^ (relative to housekeeping and control) method according to Livak and Schmittgen (2001). A gene fragment encoding alfalfa ubiquitin (UBQ, AW686873) was used as an internal control to normalize the amount of template cDNA, in agreement with the reference gene used by Kakar *et al*. (2008). The gene-specific primer pairs employed for the detection of the alfalfa transcripts of SnRK1.1 (MS.gene86051.t1), Ms.KIN_Fw: 5′CAACAGTTCCCTGCTGAGAG3′ and Ms.KIN_Rv:

5′CAACAGCCCATCTGCACTTC3′ and SENESCENCE-ASSOCIATED PROTEIN 5 (SEN5 Medtr2g079670.3) a gene with known functions in SnRK1 pathway marker genes Mt.SEN5_Fw: 5′CGTGGTCCATATCCACCAGATC3′ and Mt.SEN5_Rv:

5′GATCACAACAGATCCATCAGCTGC 3′. The design of the specific primers was performed by homology analysis between the tags of the expressed alfalfa sequences and the public database. The amplification efficiency of all primers was pre-tested, and only those that showed amplification efficiency within the range of 90 and 100% were used for qPCR. The expression analysis for each gene was repeated three times to ensure technical reliability.

## Statistical Analysis

All experiments were replicated at least three times to ensure reliability. The statistical significance of mean differences was evaluated using ANOVA, followed by Tukey’s post hoc test (p < 0.05). For pairwise comparisons, a parametric Student’s t-test was applied to identify significant changes. The complete variable metabolite dataset was used to generate the Principal Component Analysis (PCA) biplot. All data were processed and analyzed using InfoStat software (Di Rienzo *et al*., 2019).

## Results

### Reduction of leaf growth during salt stress in alfalfa

Leaf growth alterations are among the earliest responses to salt stress (Munns and Tester, 2008). In our study, we characterized the impact of salt stress on the area of the third trifoliate leaf in alfalfa plants within the experimental system (Figure 1). The leaves used in this study were identified as carbon source leaves, as they were more than 40% expanded at the onset of the treatment. We observed a significant reduction in leaf area in 3 dpt in control condition. While the leaves of control plants reached their final size by 3 dpt, the leaves of treated plants continued to grow at a slower rate compared to the initial growth period (0–1 dpt). In this regard, the ALAGR between 1 and 3 dpt was higher under control conditions (0.25 cm^2^/day), whereas in NaCl-treated plants, it was 0.04 cm^2^/day and remained at the same level until day 14. Therefore, the biochemical and molecular analysis focused on distinct stages of leaf development, providing insights into the dynamic responses of alfalfa to salt stress.

**Fig. 1.**
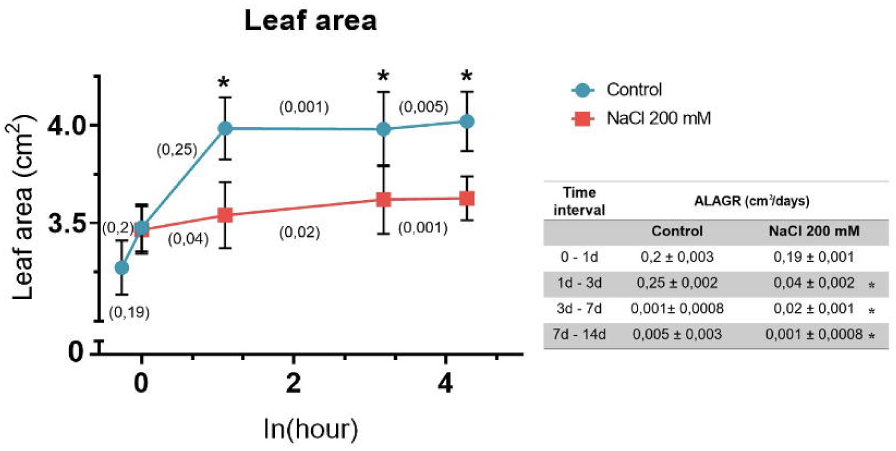
Leaf area of the third leaf of alfalfa plants after treatment with 200 mM NaCl or without (Control). Plants were grown for 20 days in 0.5X B&D medium before the treatment. Leaf area was measured over time, shown as the natural logarithm of hours (ln(hour)) to highlight early growth dynamics. Values between parentheses indicate absolute leaf area growth rate (ALAGR) expressed in cm^2^/day. The table shows ALAGR values, and each point corresponds to a mean of different plants (n = 36) of each condition and time, from three independent experiments. Asterisks indicate significant differences with controls, analyzed using ANOVA followed by Tukey’s test (p < 0.05).

### Impact of Salt Stress on chloroplast function in alfalfa leaves

In order to assess the impact of the salt treatment on chloroplast function, we measured ChlF throughout the treatment. ChlF parameters, affected by the salinity, were normalized to the control ones in each time-point (1; 7; 14 dpt) and plotted in a radar graph to depict a global view of the structure and function of PSII electron transport (Figure 2 A). Figure 2 A shows that after one day of salt treatment key indicators of ChlF, including Fv/Fm, Phi2, qL, and PI total decreased significantly. The same behavior was observed for parameters associated with chloroplast energy production and ATP synthesis, such as, gH^+^ and vH^+^, indicative of a disruption in energy metabolism pathways (Figure 2 A). Additionally, a marked increase in NPQt was observed. After 7 dpt, there were no significant changes with respect to the control treatment. However, after 14 dpt, significant declines in photosynthetic efficiency (Phi2 and qL) and chlorophyll content (SPAD) were evident. Parameters associated with the photosynthetic electron transport chain, such as Vi and Vj, exhibited a significant increase in treated plants, accompanied by a decrease in the LEF parameter. Despite these changes, the parameters related to proton gradient and flux across the thylakoid membrane (gH^+^ and vH^+^) were significantly higher in salt-treated plants (Figure 2 A).

**Fig. 2.**
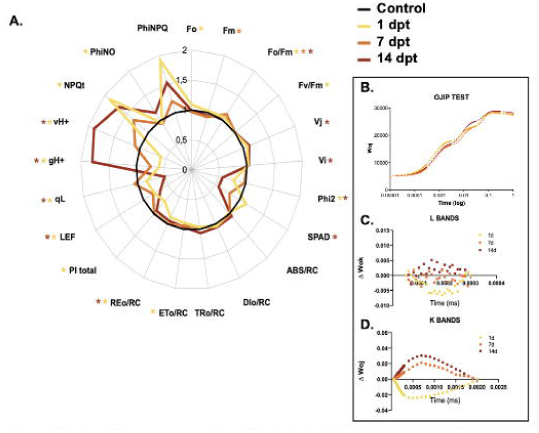
Chlorophyll fluorescence of the third leaf of alfalfa plants treated with 200 mM NaCl. A. Radar plot. B. OJIP test of control and 200 mM NaCl-treated alfalfa plants. C. K- band, fluorescence curves were normalized from the O to J steps, producing oj curves. The difference (ΔWoj) between Woj(Treated) and Woj(Control). D. L-band, normalizing fluorescence curves within 0–300 μs to create Wok curves. The difference (ΔWok), derived from Wok (Treated)- Wok(control). Different sampling times are represented by colors: Control: black; 1 dpt: yellow; 7 dpt: orange; 14 dpt: brown (dpt, days post-treatment). Asterisks of different colors indicate significant differences between sampling times under saline treatment and their respective controls. Data were obtained from four independent experiments with twenty plants per treatment at each time point (n=80) and analyzed using ANOVA followed by Tukey’s test (p < 0.05).

Analyzing the normalized ChlF curves (ΔWOK and ΔWOI) revealed previously undetectable steps, such as the K and L bands within the O–L–K–J–I–P sequence. To evaluate alterations in the OEC, fluorescence curves were normalized from the O to J steps, producing oj curves. The difference between Woj_(Treated)_ and Woj_(Control)_ highlighted K- bands (ΔWoj). Each curve in Figure 2 C and D shows the three analyzed times (1; 7 and 14 dpt). At 1 dpt, negative ΔWoj values suggested delayed electron transfer, while positive values at 7 and 14 dpt indicated OEC damage (Figure 2 C). For photosystem II (PSII) connectivity, L-band analysis involved normalizing fluorescence curves within 0–300 μs to create Wok curves. The difference (ΔWok), derived from treated minus control samples, showed minimal stress-induced disruptions (Figure 2 D).

### Temporal changes in alfalfa carbon metabolism induced by NaCl

Metabolomics serves as a critical link between plant genetics and observable traits. By analyzing the metabolome, shifts in metabolic profiles can be detected, enabling the examination of response mechanisms at the metabolic level. To analyze the dynamic changes in metabolite levels in alfalfa leaves under salt stress, leaf samples were harvested from both control and salt-stressed plants at previously described time-points and metabolite profiling was conducted using LC-MS/MS according to Figueroa *et al*. (2016). The dynamic interplay of sugar metabolism across different time points reveals a complex response to stress. Initially, within one hour of exposure (1hpt), there was an increase in Suc content and a significant decrease in Glc and ADP-Glc (Figure 3). As the stress persisted into the third hour (3hpt), Suc levels continued higher than control, alongside an increase in glucose-6-phosphate (Glc6P), + 51.5% with respect to control. Simultaneously, sucrose-6-phosphate (Suc6P) also exhibited a notable increase. Moreover, the TCA cycle was impacted by salt stress, with citrate, cis-aconitate, and isocitrate showing significant decreases in abundance under salt treatment. In contrast, 2-oxoglutarate, succinate, fumarate, and malate exhibited an opposing trend, with their levels increasing in salt-treated leaves. Moving into the first day, the TCA cycle intermediates showed the opposite than at 3 hpt, increases in citrate, cis-aconitate, and isocitrate were observed, while fumarate and malate decreased. Furthermore, there was a decrease in UDP-glucose (UDP-Glc) levels alongside a sustained increase of Suc6P. Furthermore, a decrease in fructose-1,6- bisphosphate (FBP), fructose-6-phosphate (Fru6P), and Glc6P underscored a metabolic deceleration. By the third and seventh day, there was a decrease in fructose levels coupled with a reduction in Fru6P. The TCA metabolites intermediates showed a decrease.

**Fig. 3.**
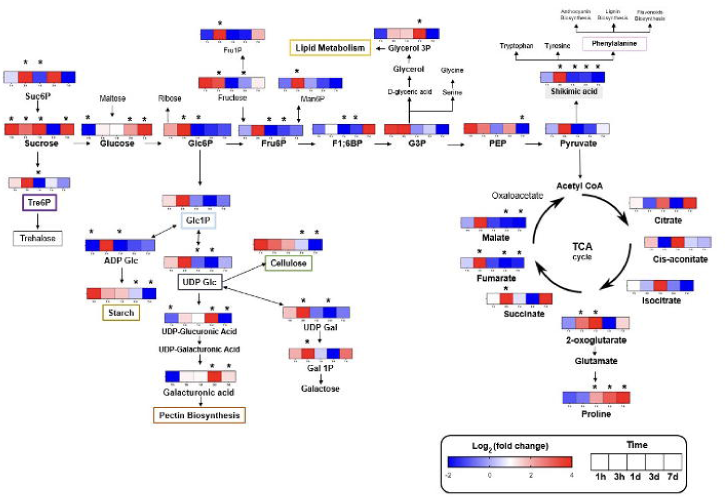
Schematic diagram of response metabolites of alfalfa under salt stress identified in this study. Heat map colors represent the log2(fold change) in the metabolite content of the stress-treated group compared to the control. Blue represents a decrease in content, white represents no change and red represents an increase in content. Each square represents a time point (1h, 3h, 1d, 3d, 7d). Asterisks indicate significant differences between plants under saline treatment and their respective controls, based on four independent experiments with four plants per treatment, grouped into one pool (n=16), analyzed using ANOVA followed by test t′Student (p< 0.05).

### Dynamic Response of Suc-Tre6P-SnRK1 to Salt Stress

The Suc–Tre6P nexus model suggests that Tre6P functions as a key signal for sucrose availability, playing a pivotal role in maintaining metabolic balance by regulating both sucrose synthesis and utilization (Yadav *et al*., 2014). Central to this model is SnRK1, which acts as an essential regulator within the signaling network (Peixoto and Baena Gonzalez, 2022). To investigate the dynamic regulation of the Suc-Tre6P-SnRK1 relationship under salt stress in alfalfa leaves, we analyzed Suc and Tre6P content along with SnRK1 activity. The SnRK1 activity was analyzed using three complementary approaches. i) using a universal kinase kit (Biosystem) to measure SnRK1 activity *in vitro*; ii) detecting the expression of a selected SnRK1 marker gene, MsSEN5 gene (*Medtr2g079670.3*), homologous to SEN5 (*AT3g15450*) in Arabidopsis, as a marker of SnRK1 activity *in vivo.* This gene has been widely employed as a sensitive indicator of SnRK1 activity in promoter activation assays and transcript abundance analysis (Baena González *et al*., 2007; Ramon *et al*., 2019). iii) Assessing the phosphorylation of the conserved T-loop domain, a hallmark of SnRK1 activation (Polge and Thomas, 2006). The expression of MsSEN5 (a readout of SnRK1 activity *in vivo*) and T-loop phosphorylation exhibited a pattern similar to that of the universal kinase during the short-term response (Figure S3 and Figure 4 C).

**Fig. 4.**
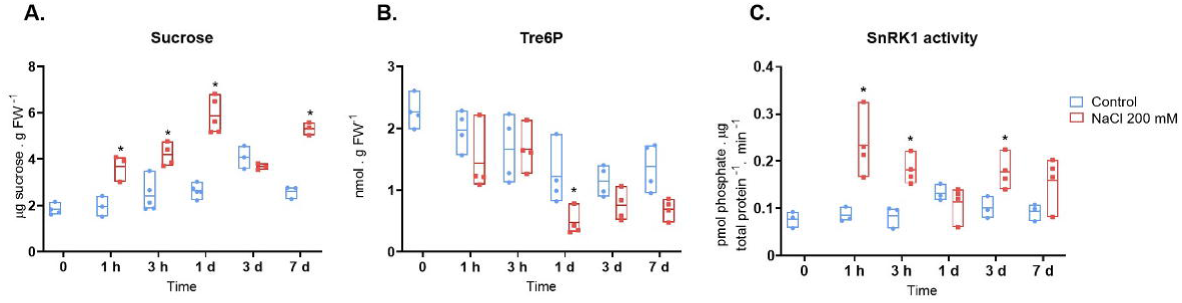
Suc-Tre6P-SnRK1 in alfalfa leaves during NaCl stress. Graphs show results for leaves from control (blue) and treated with NaCl 200 mM (red) grown for 20 days in 0.5X (Tre6P) concentration measured by LC-MS/MS. C. SnRK1 activity measured with universal kinase kit. Asterisks indicate significant differences between plants under saline treatment and their respective controls, based on four independent experiments with four plants per treatment, grouped into one pool (n=16), analyzed using ANOVA followed by Tukey’s test (p < 0.05).

In our experimental setup, the dynamic pattern of the Suc-Tre6P-SnRK1 unfolds over distinct temporal phases. Focusing on the initial response within the first hour of salt exposure (1 hpt), SnRK1 activity increased significantly in treated plants, rising by +228.6% compared to initial (0 hpt) levels and +155.5% compared to the 1 hpt control group (Figure 4 C). During this brief timeframe, Suc levels also began to increase in treated plants significantly and exhibited a negative correlation with Tre6P content (Figure 4 A-B, Figure 5 E). At 1 dpt, SnRK1 activity in treated plants began to gradually decrease (Figure 4C). This coincided with a peak in Suc accumulation and a significant decline in Tre6P levels (Figure 4 A and B). Within the first 24 h of treatment, Suc concentration and SnRK1 activity exhibited a negative correlation in salt-treated plants (Figure 5 C). Additionally, in this period, the expression of MsSEN5 and T-loop phosphorylation were also higher in salt than in control plants (Figure S3 A and B). This phase represents the broader changes occurring within the first hours of stress. By the third day (3 dpt), a long-term response was evident. SnRK1 activity rose again by +63.63% in treated plants compared to control.

**Fig. 5.**
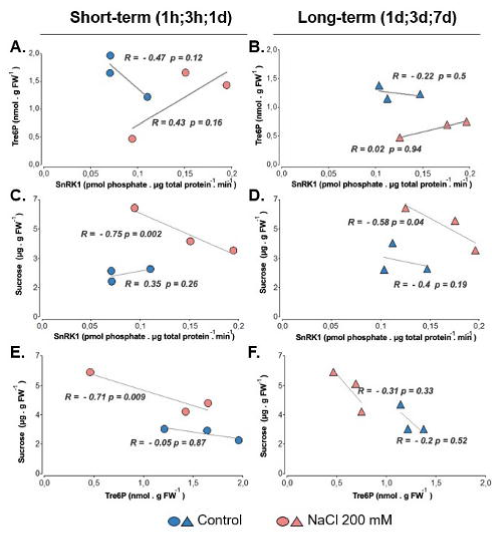
Correlations between sucrose, Tre6P, and SnRK1 under control and salt stress conditions. Linear regression analysis was performed to evaluate the relationships among sucrose, Tre6P, and SnRK1 during the short-term (1h, 3h, 1d, circles) and subsequent long- term (1d, 3d, 7d, triangles) Blue markers denote control plants, while red markers represent salt-treated plants. Both negative (R < 0) and positive (R > 0) correlations were observed.

However, transcript analysis of MsSEN5 did not follow the same trend (Figure S3 C). As stress extended to the seventh day, SnRK1 activity tended to decline once again, while a significant increase in Suc concentration was observed. During this prolonged phase, a negative correlation between Suc concentration and SnRK1 activity was observed (Figure 5 D).

### Effect of Sucrose and NaCl on SnRK1 Activity

Since we observed that SnRK1 and Suc exhibited contrasting trends during salt stress (Figure 4 and 5), we questioned whether the increase in sucrose induced by salt stress could be influencing SnRK1 activity. To explore this, we supplied Suc at physiological levels (10 mM), using darkness as a positive control (SnRK1 activator) and light as a basal control (Figure 6). Additionally, we included treatments with NaCl (100 mM) and a combination of NaCl (100 mM) + Suc (10 mM). In this experimental system, the 100 mM NaCl treatment was applied directly to the leaf, unlike the 200 mM root-fed NaCl treatment used to analyze leaf responses. Soluble sugars, SnRK1 activity and Na^+^ concentration were measured in all samples (Figure 6 B and Figure S4).

**Fig. 6.**
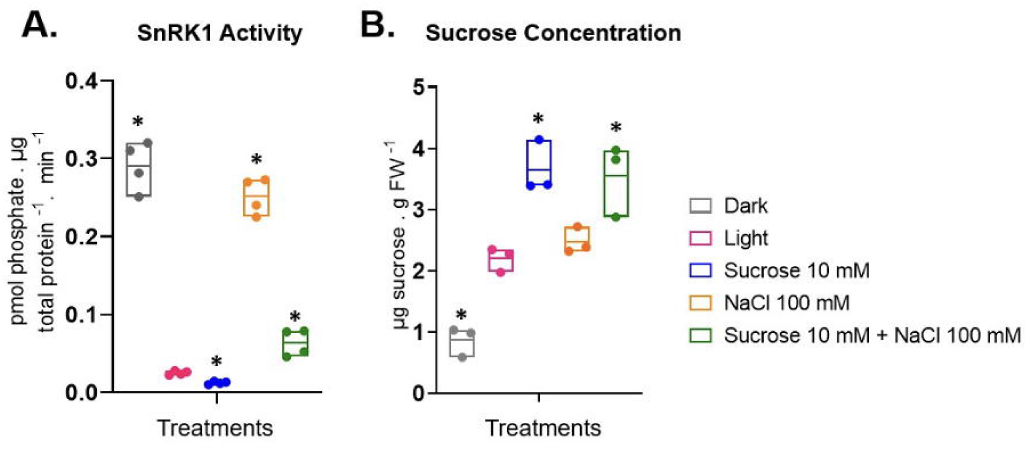
Effect of Sucrose and NaCl on A. SnRK1 activity and B. sucrose concentration in alfalfa leaves cultured for 20 days in B&D medium 0.5X and subsequently subjected to different treatments during 3 h: darkness (positive control); light (basal control); sucrose 10 mM; NaCl 100 mM; combination of sucrose 10 mM and NaCl 100 mM. Asterisks indicate significant differences of treated leaves compared to the control (light) in three independent experiments with four plants per treatment, grouped into one pool (n=12), analyzed using ANOVA followed by Tukey’s test (p < 0.05).

Alfalfa leaves were incubated for 3h with 10 mM Suc, resulting in a significant decrease in SnRK1 activity (-50.9% relative to the light control). To investigate whether this effect persisted under stress conditions, we conducted further experiments. When leaves were exposed to a 3-hour treatment with 100 mM NaCl, SnRK1 activity exhibited a significant increase (+155% relative to the light control) without changes in Suc content (Figure 6 A and B). However, when Suc was applied in combination with 100 mM NaCl, SnRK1 activity displayed a decrease (-74.8% relative to the 100 mM NaCl treatment alone). Na^+^ concentration was measured and showed a significant increase in leaves treated with 100 mM NaCl compared to light control (Figure S4 A).

### Temporal associations of SnRK1 and metabolites in salt-stressed leaves

To gain a comprehensive understanding of our results and evaluate the effect of time on salt-stressed and control alfalfa leaves, a principal component analysis (PCA) was performed independently for salt-treated and control plants (Figure 7 A and B). These analyses explained 78.4 % and 69.4 % of total variation in salt-treated and controls, respectively (Figure 7). The analysis revealed distinct temporal associations of SnRK1 and metabolites. In salt-stressed leaves, SnRK1 activity was linked to the early response (1 hpt), whereas metabolites associated with anabolic processes, such as Tre6P or shikimate, were predominantly observed at 3 hpt (Figure 7A). Furthermore, intermediates of the TCA cycle, including isocitrate, fumarate, and malate, were also associated with this time point. Metabolites related to cell-wall composition were primarily linked to later stages, at 3 dpt.

**Fig. 7.**
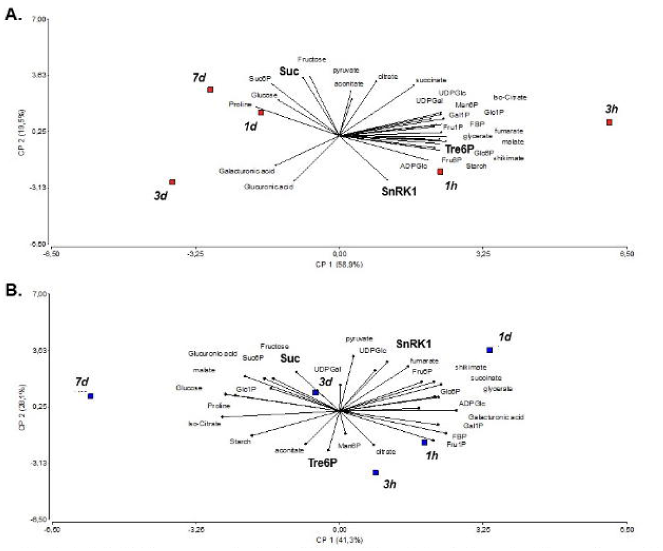
Principal Component Analysis (PCA) biplot obtained from the first and second principal components A. Leaves of NaCl 200 mM treated plants; B. control leaves. Measurements were made on the third trifoliate leaf of alfalfa plants. Results are from four independent experiments with a pool of three leaves per experiment.

In control leaves, a distinct pattern was observed. Tre6P remained associated with 3 hpt, while Suc was linked to 3 dpt. SnRK1, in contrast, showed a stronger association with 1 dpt (Figure 7B).

## Discussion

Alfalfa is one of the most widely cultivated forage crops, valued for its adaptability and high nutritional content. However, salinity poses a significant challenge to its productivity, particularly in arid and semi-arid regions. After 14 days of hydroponic salt treatment (200 mM NaCl), Kumen alfalfa plants showed significant growth reductions compared to controls, including 37.4% in height, 38.4% in leaf number, and 50% in aboveground dry weight (Barbieri *et al*., 2024). Our study demonstrated that salinity during alfalfa’s vegetative phase impairs leaf growth, alters chloroplast function, and induces metabolic shifts. Meta-analyses have revealed that leaf-specific area and leaf dry mass per unit area are among the earliest parameters affected by salinity (Munns and Tester, 2008; Poorter *et al*., 2010). Leaf area expansion was arrested as early as 1 dpt under salt stress, with an ALAGR of 0.04 cm^2^/day between 1 and 3 dpt, compared to 0.25 cm^2^/day in the control during the same period (Figure 1). For optimal growth and development, the Na^+^/K^+^ ratio in the cytosol of plant cells typically ranges from 1–10 mM Na^+^/100–200 mM K^+^ (Kader *et al*., 2006). In our study, this ratio increased significantly after 1 hpt (+177%) (Figure S2 C). Noticeably, the Na^+^ concentration increase at 1 hpt may reflect vacuolar compartmentalization rather than cytotoxicity.

Salt stress disrupts multiple structures and processes in plants, among them the photosynthetic apparatus and stomatal conductance. These effects have been well-documented in various plant species, such as rice and barley (Munns and Tester, 2008; Acosta-Motos *et al*., 2017). Photosynthetic efficiency (PI), a key indicator of plant responses to abiotic stress, exhibited a transient decline at 1 dpt (Figure 2A). Similarly, the maximum quantum yield of PSII (Fv/Fm), which reflects PSII integrity, showed a significant but temporary reduction at the same point. PSI electron acceptor side (REo/RC) showed a significant decline from 1 dpt. Accordingly, Kaiwen *et al*. (2020) reported that PSI in alfalfa leaves was more sensitive to salt stress than PSII. Chlorosis symptoms in salt-treated alfalfa leaves were quantified using the SPAD index, which showed a 40% reduction at 14 dpt compared to untreated plants (Figure 2 A). At the same time, salt stress disrupted the activity of the OEC on the donor side of PSII and impaired electron transport to the secondary quinone (QB) on the acceptor side suggested by an increase in Vj (Figure 2). In alfalfa, the PSII acceptor side exhibited greater sensitivity to salt stress compared to the donor side (Li *et al*., 2024). Rises in Vi and the K-band are linked to damage in the OEC on the donor side of PSII (Strasser *et al*., 2004). Consistent with these findings, Li *et al*. (2020) reported that alfalfa seedlings exposed to 200 mM NaCl for 14 days showed significant reductions in photosynthetic rates and disruptions in Rubisco activity. Overall, the damage to chloroplast function and imbalances in the electron transport chain caused by salt stress may lead to metabolic alterations and impaired plant performance.

Sugars in plants fluctuate significantly in response to photosynthetic activity and environmental variables. Our results showed an increase in Suc associated with an alteration of the photosynthetic apparatus (Figures 2 and 3). In tomato, salt stress decreases Suc transport from leaves to roots, leading to carbohydrate accumulation in the leaves (Suwa *et al*., 2008). Furthermore, reduced sink demand under these conditions results in elevated Suc levels in photosynthetic tissues, which in turn contribute to feedback inhibition of the Calvin-Benson cycle (Flowers *et al*., 2009). Additionally, a decreased utilization of carbon skeletons in source leaves for growth or energy production could be related to high sugar accumulation. In this regard, sugar accumulation did not promote growth or high leaf growth rates (Figures 1 and 3). Moreover, sugar accumulation might be related to counteract osmotic stress. This mechanism was demonstrated in salt-tolerant cultivars, where the fine-tuned regulation of starch and sucrose metabolism promotes the accumulation of soluble sugars such as sucrose, maltose, glucose, and trehalose (Gao *et al*., 2018).

Most studies on alfalfa stress responses focus on long-term transcriptional effects, often overlooking short-term mechanisms and post-translational signaling (Wang *et al*., 2021; 2024). Here, we evaluated metabolite profiling in salt-treated plants at both short- and long-term treatment (Figure 3). Previous research by other authors had revealed distinctive patterns of metabolites (D’Amelia *et al*., 2018; Derakhshani *et al*., 2020). For example, salt-stressed wheat exhibits significant metabolic changes, including increased levels of Suc and amino acids like proline, alongside reductions in organic acids and TCA cycle intermediates (Che-Othman *et al*., 2019). In our study, we observed a notable metabolic shift at 3 hpt, marked by decreases in citrate, aconitate, and isocitrate accompanied by an increase of 2-OG, succinate, fumarate and malate (Figure 3). These changes suggest disruptions in the TCA cycle, potentially driven by an earlier upregulation of SnRK1, a critical regulator of energy balance during stress (Figure 3 and 4 C). Supporting this hypothesis, plants with reduced SnRK1 activity had elevated levels of citrate, aconitate, and isocitrate, accompanied by lower levels of 2-OG and succinate indicating altered flux of metabolites into the TCA cycle (Peixoto *et al*., 2021). In our work, the TCA cycle metabolites shifts may result from salt-induced inhibition of key enzymes, such as pyruvate dehydrogenase, which converts pyruvate to acetyl-CoA. This inhibition is supported by the observed significant accumulation of pyruvate (Figure 3). Similarly, 2-OG dehydrogenase, which facilitates the conversion of 2-OG to succinyl-CoA, may also be negatively affected under salt stress (Che-Othman *et al*., 2019). To compensate for these enzymatic alterations, plants may activate the GABA shunt, providing an alternative pathway for carbon flow to sustain respiration during stress (Che-Othman *et al*., 2019). Noticeably, Li *et al*. (2020) reported a marked increase in TCA cycle metabolites in alfalfa seedling leaves treated with 200 mM NaCl for 14 days. The apparent discrepancies between these findings and our results may stem from the timing of measurements. Our data highlights that the plant’s metabolic response to stress is dynamic, with fluctuations in metabolite levels—both increases and decreases—occurring over the period of stress exposure. Moreover, principal component analysis (PCA) further supports this dynamic response, revealing temporal associations of metabolites under salt stress, with TCA cycle metabolites predominating at 3 hpt, and later cell-wall-related changes emerging at 3 dpt (Figure 7 A).

The significant accumulation of Glc6P at 3 hpt (+51.5% relative to the control) points to the activation of hexokinase (HXK). Rather than supporting glycolysis as a primary energy source, this response is likely to facilitate the oxidative pentose phosphate pathway (OPPP), which provides reducing power (NADPH) and metabolic intermediates crucial for defense response, such as the shikimate pathway. Phenylalanine, an intermediate of the shikimate pathway, is closely linked to stress-defense mechanisms. By 1 dpt, reduced levels of shikimate may signal growth arrest through the suppression of key metabolic regulatory proteins (Bledsoe *et al*., 2017). The consistently lower shikimate levels in salt-treated plants suggest a redirection of energy. This shift moves reserves, such as starch and sugars, away from growth-related activities like amino acid biosynthesis. These findings agree with similar metabolic responses reported in salt-stressed Arabidopsis, *Lotus japonicus*, and rice (Sanchez *et al*., 2007).

The metabolic shifts detected at 3 hpt and the use of alternative energy sources may be linked to earlier signaling events. SnRK1 is a crucial regulator of Suc homeostasis, playing a role in plants analogous to its mammalian counterpart’s function in glucose regulation (Peixoto and Baena Gonzalez 2022). The rapid activation of SnRK1 observed within the first hour of salt stress (+228.57%) suggests its involvement in subsequent Suc accumulation (Figure 4). Accordingly, plants SnRK1α-OE displayed elevated sucrose (Suc) levels (Li and Zhao, 2024). However, since SnRK1 activation can also inhibit sucrose phosphate synthase (SPS) activity, it may simultaneously limit Suc synthesis (Peixoto and Baena Gonzalez, 2022). This apparent contradiction may result from enhanced Suc synthase activity and broader metabolic adjustments. Furthermore, a negative correlation between SnRK1 activity and Suc content during the short-term response (-0.75; 1 hpt to 1 dpt) suggests that Suc may act as an inhibitor of SnRK1 activity (Figure 5). Experimental evidence demonstrated that Suc acts as a negative regulator of SnRK1 activity under both basal and salt stress conditions (Figure 6). According to this, the expression of SnRK1 marker genes were significantly reduced in sugar-treated Arabidopsis seedlings as compared to the control (Pereyra *et al*., 2023). In contrast, in salt-stressed alfalfa leaves, 100 mM NaCl treatment significantly increased SnRK1 activity (+155%), but when combined with 10 mM Suc, this activation was markedly suppressed (-74.8% relative to NaCl 100 mM). Suc seems to act as a negative modulator of SnRK1 activity; however, other signaling pathways, such as redox state and hormonal signaling, likely contribute to SnRK1 activation during NaCl stress. In addition, the observed increase in Suc concentration may contribute to osmotic balance, cellular integrity, and the activation of stress-signaling pathways (Ruan 2014). However, whether this accumulation results from reduced utilization, enhanced synthesis, or a combination of both remains an open question. Tre6P is a well-established regulatory molecule essential for growth, with significant impacts on metabolism and development (Fichtner and Lunn, 2021). According to the Tre6P-sucrose nexus model, Tre6P signals Suc availability, maintaining homeostasis by regulating Suc production and utilization (Yadav *et al*., 2014). Elevated Tre6P-to-sucrose ratios in actively growing tissues enhance Suc utilization and import (Yadav *et al*., 2014; Fichtner and Lunn, 2021). In salt-stressed alfalfa leaves, however, Tre6P-to-Suc ratio was significantly disrupted (Figure 4). Suc and Tre6P levels exhibited a strong negative correlation (R= -0.71 from 1 hpt to 1 dpt), with Suc accumulating (Figure 5). The negative correlation between Tre6P and sucrose may result from their distinct subcellular localization. Tre6P, synthesized in the cytosol, reflects nucleo-cytosolic sugar levels, which can be masked by high vacuolar sugar concentrations in alfalfa leaves cells. Moreover, its production is especially prominent in guard cells and near the phloem-loading zones of source leaves—key regions for regulating source-sink dynamics and systemic signaling (Fichtner *et al*, 2020). The sucrose–T6P nexus may adjust during alfalfa development to match changing metabolic needs. Furthermore, Tre6P levels did not correlate with SnRK1 activity, reflecting a breakdown in homeostasis (Figure 4). At 1 dpt, Suc levels peaked while Tre6P significantly declined, despite SnRK1 activity returning to baseline levels. In Arabidopsis mature leaves, the SnRK1 inhibition by Tre6P is weaker than in growing tissues (Zhang *et al*., 2009). Tre6P levels were still associated with the expression of SnRK1 marker genes (Peixoto *et al*., 2021; Avidan *et al*., 2023). Other interacting components, such as upstream protein kinases (Zhai *et al*., 2018) and class II TPS proteins (Van Leene *et al*., 2022), may influence SnRK1’s response to T6P (Zhang *et al*., 2009). However, the actual mechanisms by which these interactions function in alfalfa mature leaves under salt stress remain unclear. Low Tre6P, combined with active SnRK1, likely promotes stress responses and catabolism over anabolic processes (Baena González *et al*., 2007; Zhang *et al*., 2009). Overexpression of SnRK1 (SnRK1-OE) has been associated with "Tre6P hyposensitivity" which partially blocks the feedback effects of Tre6P on Suc metabolism (Peixoto *et al*., 2021). Supporting this, Peixoto *et al*. (2021) demonstrated that plants SnRK1-OE had higher Tre6P:Suc ratios, while SnRK1 mutants displayed lower ratios. This highlights SnRK1’s role in balancing Suc and Tre6P levels. However, salt-treated alfalfa showed an increase in SnRK1 with a significant decrease in Tre6P:Suc ratio at 1 hpt. The disrupted Tre6P:Suc ratio indicates a decoupling of the sucrose–Tre6P regulatory nexus under salt stress. According to the PCA, both SnRK1 and Tre6P play a significant role during the short-term response to salt stress, whereas sucrose appears to be involved in later responses, potentially linked to changes in the cell wall. Nunes *et al*. (2013) demonstrated a strong interplay between Tre6P, Suc, and SnRK1-regulated gene expression, independent of growth rate. They proposed that Tre6P is not a direct growth signal but, through SnRK1, primes gene expression to support growth. The model establishes that Tre6P mediates changes in gene expression via SnRK1, aligning growth with Suc supply. Altogether, these findings highlight the complexity of sugar metabolism during stress responses and the need for further exploration of regulatory mechanisms.

In this study, we propose that early biochemical shifts are closely linked to subsequent stress responses, identifying the rapid activation of SnRK1 as a key determinant of salt stress tolerance in alfalfa—an insight that opens new avenues for future research. Notably, we provide the first evidence of a SnRK1 wave-like activation pattern over time and a disruption in the Tre6P-Suc regulatory link in alfalfa. Together, these alterations underscore the crucial role of early SnRK1 activity in maintaining chloroplast function and driving the metabolic reprogramming necessary for energy production and fine-tuned TCA cycle regulation. These findings provide a base for advancing our understanding of the complex metabolic networks that govern plant stress tolerance.

## Supplementary data

Fig. S1. Experimental set-up

Fig. S2. Ion concentrations in leaves of alfalfa

Fig. S3. Effect of salinity on AMPK phosphorylation and SnRK1 marker gene expression in alfalfa leaves.

Fig. S4. Effect of Sucrose and NaCl on Na^+^, glucose and fructose content

Table S1. Description of ChlF parameters

Table S2. Metabolite ratios in alfalfa leaves

## Acknowledgments

We thank Monica Cornachione and Ariel Odorizzi for providing alfalfa seeds and Technician Paola Suarez for technical assistance. Prof John Lunn for metabolomic analysis collaboration. Prof Edith Taleisnik for a critical review of this manuscript.

## Author contribution

MR conceptualization. GB and MR designed and performed the experiments. GB, RP methodology. RF performed the LC-MS/MS measurements and analyzed the data. GB, RP and MR formal analysis. GB and MR created and edited the final Figures. GB, RP, MR writing - original draft. GB, RP, RF, MR writing - review & editing. MR: supervision; MR: funding acquisition.

## Data availability

The data that support the findings of this study are available from the corresponding author upon reasonable request.

## Abbreviations

2OG,: 2-Oxoglutarate;
ChlF,: Chlorophyll fluorescence;
dpt,: Day post treatment;
hpt,: Hour post treatment;
OEC,: oxygen-evolving complex;
PCA,: Principal component analysis;
SnRK1,: Sucrose non-fermenting kinase 1;
TCA,: Tricarboxylic acid;
Tre6P,: Trehalose 6- phosphate.

---

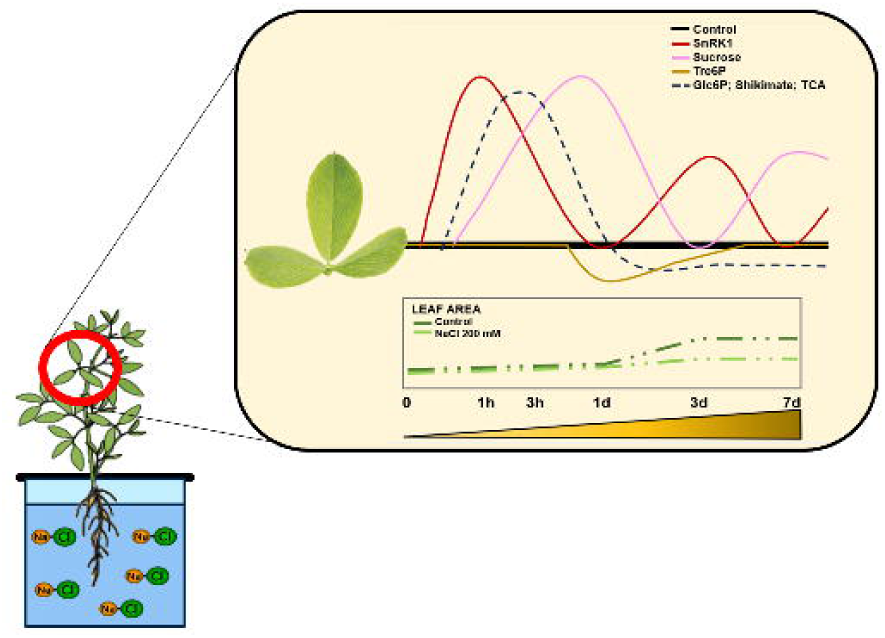

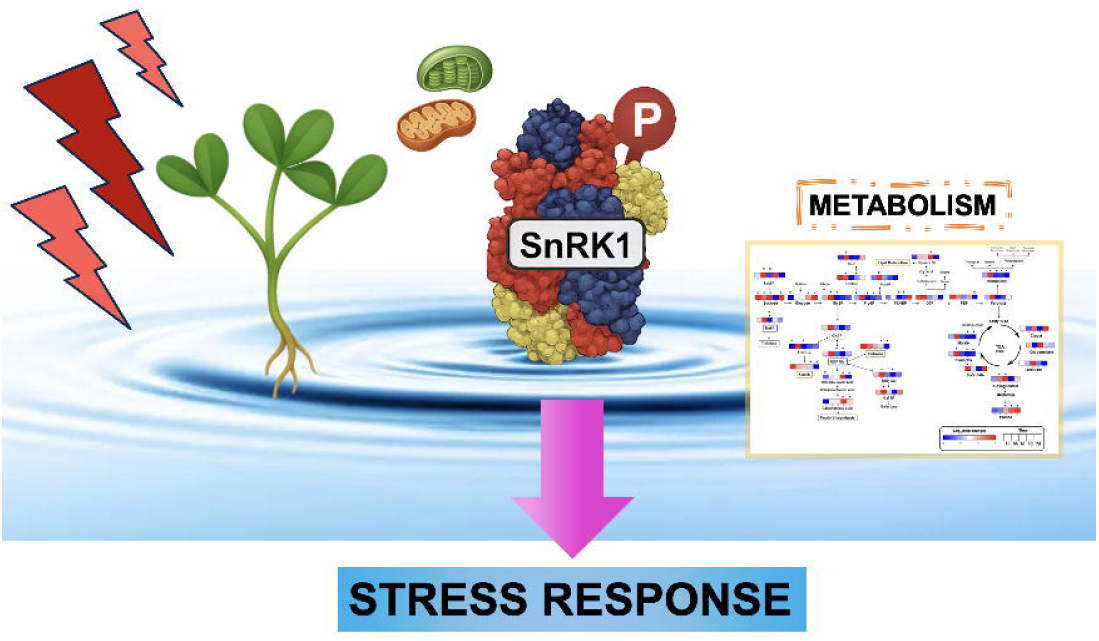

